# Pause characteristics of sentence production in Parkinson’s disease: insights from sentence complexity and length

**DOI:** 10.1101/2025.08.07.669233

**Authors:** Fatemeh Mollaei, Huw Evans, Alexandra Pool

## Abstract

**Purpose:** Parkinson’s disease (PD) affects forward flow of speech including fluency disruptions in 90% of individuals. One of the main parameters affecting flow and fluency of speech is pause behaviour. However, the precise language characteristics of pauses, including sentence complexity and length, and how they contribute to the fluency disruptions of PD are not fully understood. This study examined how sentence complexity and length affect pause behaviour in PD.

**Method:** Seventy-one participants, comprising individuals with PD (n = 32) and neurotypical controls (n = 39), read a speech passage aloud. The number and duration of pauses, categorised by location (between, within sentences), sentence complexity (simple, complex), and sentence length (short, long) were analysed. Cognitive ability, assessed using the Montreal Cognitive Assessment (MoCA), and motor speech deficits (i.e., dysarthria) severity, assessed using a speech perceptual ranking, were evaluated and correlated with pause characteristics.

**Results:** Individuals with PD produced significantly more pauses across all categories compared to controls. However, only between-sentence and long-sentence pauses were significantly longer in duration. Pause frequency and duration in both groups were higher in more complex and longer sentences. Significant negative correlations were found between MoCA scores and number of pauses. Significant positive correlations were observed between dysarthria severity and duration of pauses.

**Conclusion:** These findings suggest that increased cognitive-linguistic demands—indexed by sentence complexity and length—may underlie pausing behaviour and contribute to fluency disruptions in individuals with PD. The results extend previous research by highlighting the potential cognitive-linguistic basis of motor speech dysfunction in PD.

## Introduction

Neurodegenerative disorders affect millions of people worldwide, with Parkinson’s disease (PD) representing the fastest growing and progressively debilitating condition. PD affects approximately 2% of individuals over the age of 65 [1] and is marked by the degeneration of dopaminergic neurons in the substantia nigra [2]. This dopamine loss disrupts basal ganglia function and impairs motor coordination, leading to hallmark symptoms such as bradykinesia, rigidity, postural instability, tremor, and diminished respiratory support [3]. As no cure currently exists, treatment primarily targets symptoms management and the slowing of disease progression [4].

These motor symptoms often extend to the speech mechanism, resulting in hypokinetic dysarthria, a motor speech disorder characterized by reduced loudness, monopitch, monoloudness, breathy and hoarse vocal quality, and imprecise articulation [5]. Such deficits can compromise communication and negatively impact quality of life. Despite clinical guidelines recommending multidisciplinary management, including speech-language therapy (SLT) [6], only a small proportion (3–4%) of individuals with PD receive SLT services, even though up to 90% develop speech impairments [7]. This underutilization likely reflects a range of systemic barriers, including limited resources, late-stage referrals, and a lack of standardized criteria for prioritizing SLT care [8,9]. Identifying objective, speech-based markers of impairment could help guide SLT referrals and treatment decisions.

### Speech impairments in Parkinson’s disease

Speech in PD is most commonly impaired by hypokinetic dysarthria, which affects phonation, articulation, and prosody. Vocal deficits often stem from vocal fold bowing and incomplete glottal closure [10], while articulation is marked by reduced movement amplitude and velocity [11]. Prosodic deficits include monotone pitch and loudness, as well as atypical speech rate and pausing patterns [12]. These disruptions, coupled with reduced facial expressiveness and gesture control, contribute to decreased speech intelligibility— particularly in later stages of the disease [13]. Although the precise relationship between disease severity and motor speech impairment remains inconclusive [14], most research suggests a progressive decline in speech intelligibility as PD advances [15].

Speech-language interventions often target respiratory support and speech rate modulation to improve overall fluency [16]. However, a comprehensive understanding of how PD affects specific components of speech—such as pause behaviour—remains limited.

### Fluency and pause behaviour in PD

Speech fluency involves the smooth, continuous production of speech, and is shaped by the timing of pauses and modulation of speech rate [17]. In PD, dysfluencies are considered neurogenic, arising from neurological disruption rather than developmental [18]. While phonation and articulation in PD have been extensively studied, less attention has been paid to fluency and pause timing—despite consistent clinical observations of rapid, irregular speech interspersed with atypical pauses [19].

Articulatory findings in PD are mixed. Some studies report slower syllable repetition rates due to diminished tongue strength and speed [20], as well as impaired control of sentence duration [21]. Others report increased articulation rates, potentially linked to impaired motor timing [22,23]. Regardless of articulation rate, pause time was found to be a reliable differentiator between individuals with PD and healthy controls [24].

### Silent pauses and cognitive influence

Although crucial to fluent speech, the role of silent pauses in PD remains underexplored. Early studies offered conflicting results—some found no differences in pause behaviour [25], while others observed more frequent and longer pauses, especially at sentence onsets [26,27]. Silent pauses in PD may reflect more than just motor dysfunction; they may also indicate underlying cognitive-linguistic challenges, such as deficits in lexical retrieval and sentence planning [28].

In healthy individuals, pauses are often placed before complex syntactic structures, where cognitive demand is greatest [29,30]. Older adults also pause more frequently, likely due to declines in working memory and processing efficiency [31]. PD is associated with early cognitive decline, particularly in executive functioning, which includes working memory, planning, and inhibitory control [32,33]. These cognitive deficits likely contribute to dysfluent speech and atypical pausing patterns [34,35].

### Sentence complexity and length in relation to pause behaviour

Few studies have examined how sentence complexity and length influence pause behaviour in PD. In typical speakers, longer and more complex sentences are associated with increased pre-boundary and intra-sentence pauses [36,37]. These pauses may help manage the greater cognitive-linguistic load associated with complex syntax [38]. It remains unclear whether individuals with PD, who often have both cognitive and respiratory impairments, show increased sensitivity to sentence complexity and length in their pause behaviour.

### Aims and research questions

This study aims to clarify how sentence complexity and length influence speech pausing in individuals with PD. We examined whether PD speakers produce more frequent and longer-duration silent pauses than neurologically healthy controls, and whether these patterns vary based on sentence complexity and length. We also investigated how cognitive function and disease severity interact with linguistic variables in shaping pause behaviour.

Three research questions were addressed:

1. Do speakers with PD produce a greater number and longer duration of silent pauses than control speakers?
2. Does sentence complexity and length affect pause behaviour?
3. How do cognitive ability and dysarthria severity interact with sentence complexity and length to influence pause frequency and duration?

We hypothesized that individuals with PD would produce a higher number of pauses and longer average pause durations compared to controls. We also anticipated that more complex and longer sentences would elicit more frequent and longer pauses in the PD group. Finally, we expected that cognitive ability and dysarthria severity would modulate the effects of sentence complexity and length on pause behaviour. Findings from this study aim to inform SLT assessment and management strategies by identifying linguistic and cognitive markers of fluency disruptions in speech in PD.

## Materials and methods

### Participants

A total of 71 participants were included in the study, divided into two groups: 39 older control participants (OC) and 32 participants diagnosed with idiopathic Parkinson’s Disease (PD). The PD group comprised of 23 males (71.88%) and 9 females (28.13%) with a mean age of 67.90 years (SD = 6.47). The mean disease duration from diagnosis was 73.41 months (SD = 62.14), ranging from 3 months to 19 years and 3 months. The OC group consisted of 39 participants, age- and gender-matched to the PD group, with a mean age of 62.71 years (SD = 8.16). Biographical information of participant groups is presented in Table 1. For demographic details please refer to the S1 and S2 Tables.

**Table 1.**
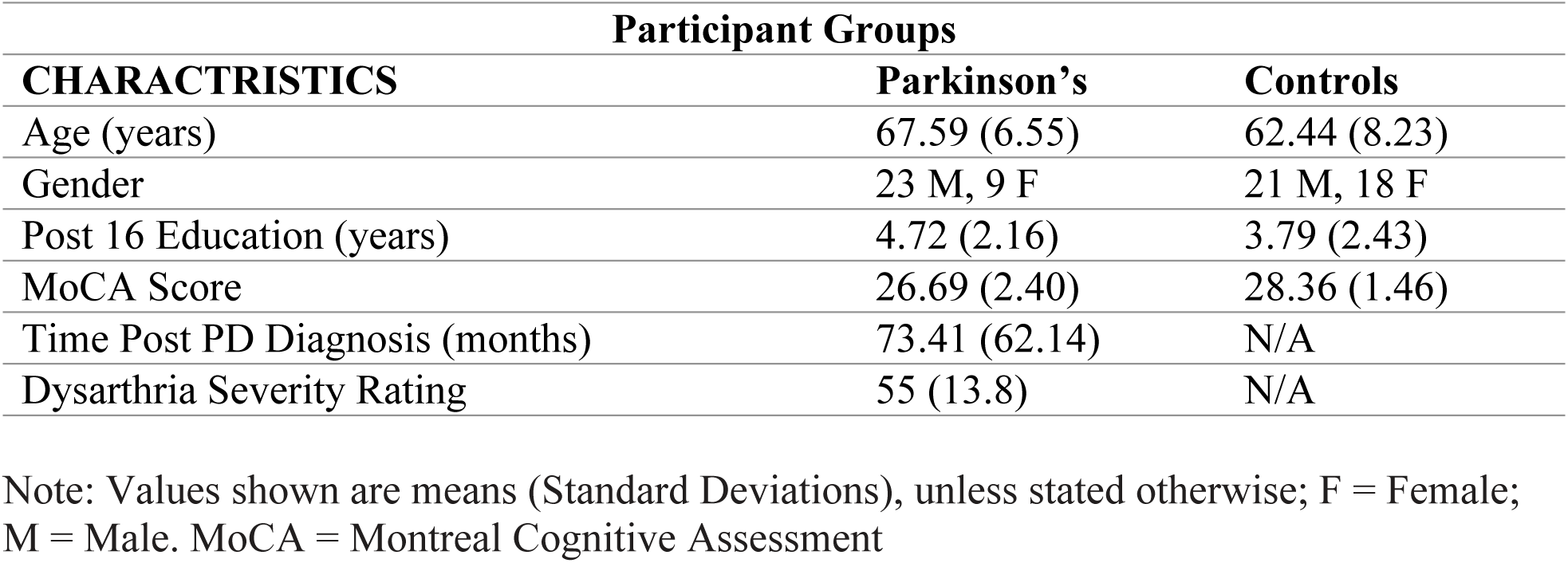
Biographical characteristics of participant groups.

The data for this study was obtained from pre-collected datasets (PD 1-18) from Mollaei et al. [39] and 46 additional data from two final-year Speech and Language Therapy students at the University of Reading (UoR). Ethical approval was granted by the UoR Research Ethics Committee and the McGill Faculty of Medicine Institutional Review Board. Prior to participation, all individuals provided fully informed written consent, including consent for the anonymized use of their data in subsequent studies. The study recruitment period took place between 10 June 2021 to 7 March 2023.

All participants were instructed to refrain from taking PD medication for at least 12 hours before data collection. Additionally, no participants had other medical or neurological conditions, including uncorrected hearing or visual impairments.

### Study design

A cross-sectional mixed-method design was used. For between-group comparisons, the independent variable was disease status, with two levels: PD and OC. For within-group analyses, the independent variable was sentence type, which included five levels: within-sentence silent pauses (SP) for short sentences, long sentences, simple sentences, complex sentences, and between-sentence silent pauses. The categorization of sentence types and passage sentences is presented in Appendices 1 and 2 based on DeDe and Salis [40]. The dependent variables were as follows:

1. Number of pauses per sentence type
2. Duration of within-sentence SP (seconds)
3. Duration of between-sentence SP (seconds)

### Procedure

Each participant completed the Montreal Cognitive Assessment (MoCA) as a short cognitive screen [41]. The MoCA was delivered in person by Mollaei et al. [39], and virtually during the UoR data collection due to COVID-19 adaptations. The standard “Rainbow Passage” [42], a controlled speech sample of 327 words, was provided to participants. In Mollaei et al.’s study, the passage was presented in person, while during the UoR collection, the passage was sent electronically in advance. Participants had the option to print it out or read it on a computer screen.

In Mollaei et al. [39], the passage was read aloud by participants and recorded for later analysis. During the UoR data collection, recordings were captured via Zoom (Zoom Video Communications Inc., 2016). Participants were instructed to sit in a quiet room with minimal distractions, and headphones/earphones were used to enhance sound quality. Task instructions were sent in advance, and participants were asked not to read the passage prior to the session to avoid adaptation effects [43].

At the beginning of each session, the study’s purpose and aims were explained, and participants had the opportunity to ask questions. Demographic information was collected using screening questions. The MoCA was administered during the session. Participants were then asked to read the “Rainbow Passage” aloud as they normally would. Audio recordings were taken for analysis.

Audio recordings were saved in WAV format and imported into Praat software (version 6.1.04) [44]. The audio files were visually displayed in Praat, allowing coders to view and listen to the recordings simultaneously. A TextGrid was created in Praat to mark silent pauses, defined as intervals exceeding 150 ms. For this study, pauses were set at a boundary of 200 ms, based on recommendations by DeDe and Salis [40], to capture shorter pauses while excluding those that likely corresponded to breath or articulation.

The coding system included categories such as silent and filled pauses within-sentence, between-sentence silent and filled pauses, false starts, mazes, revisions, and repetitions (S3 Table provides definitions) based on previous studies [40]. Breaths were treated as silent pauses due to the inability to distinguish individual breath instances. Each sentence was coded individually in the TextGrid to ensure accurate pause placement.

The coded data was exported to Microsoft Excel, and the following measures were extracted for analysis:

- Mean duration of between-sentence pauses
- Total number of pauses
- Mean number of pauses per sentence type (short, long, simple, complex)
- Mean duration of pauses per sentence type (short, long, simple, complex)

Passage recordings were additionally analyzed for clinical dysarthria severity, rated by a trained Speech and Language Therapist (SLT) using a 7-point scale (1 = normal speech, 7 = severe speech) based on perceptual characteristics related to articulation, resonance, prosody, phonation, and respiration [45].

Inter-rater reliability was assessed using qualitative ratings for the number of pauses and duration of pauses. The reliability coefficients were 0.93 for the number of pauses and 0.86 for pause duration, indicating excellent agreement between raters [46].

### Statistical analysis

Statistical analyses were conducted using SPSS software, version 28 (IBM Corp., 2021). Descriptive statistics were generated to examine the number and duration of pauses in all sentence types, separately for the PD and OC groups. Descriptive statistics for between-sentence pauses were also computed.

Due to the presence of extreme outliers in some sentence types, these were removed from the analysis: OC25, PD20, and PD27. After removing the outliers, descriptive statistics were recalculated. While the PD group did not meet normality assumptions, the OC group did, as indicated by normality tests (S4 Table). To accommodate for this violation, parametric repeated measures ANOVAs were conducted, which are robust to non-normality in large samples [47]. Disease status (PD vs. OC) was the independent variable, and the dependent variables were the duration of pauses and the number of pauses, compared across sentence complexity (simple vs. complex) and sentence length (short vs. long).

To explore the effects further, independent-sample t-tests were performed and corrected for multiple comparisons using the Bonferroni correction. For the 10 tests, the Bonferroni-corrected alpha-level was set at 0.005. Paired-sample t-tests were conducted to examine within-subject effects, with the corrected alpha-level for the PD and OC groups set at 0.0125.

Finally, Spearman’s correlations were performed to examine the relationships between dysarthria severity and cognitive ability, as measured by the MoCA and dysarthria severity scores, and the number and duration of pauses at different sentence levels within the PD group.

## Results

### Research Question 1: Do speakers with PD produce a greater number and longer duration of silent pauses than control speakers?

A significant main effect of disease status was found for the number of pauses produced in short and long sentences (*F*(1, 66) = 11.905, *p* < 0.001), with PD group (mean = 2.185) producing more pauses than the older control (OC) group (mean = 1.595). Additionally, a significant effect was found for sentence length (*F*(1, 66) = 327.827, *p* < 0.001), with long sentences (mean = 2.676) exhibiting more pauses than short sentences (mean = 1.033). No significant interaction was observed between the mean number of pauses across sentence length and disease status (*F*(1, 66) = 3.938, *p* = 0.051), although the result was marginally significant. Fig 1 illustrates that the PD group exhibited a higher number of pauses compared to the OC group across both sentence types. It also demonstrates that longer sentences had more pauses than shorter sentences in both groups.

**Fig 1.**
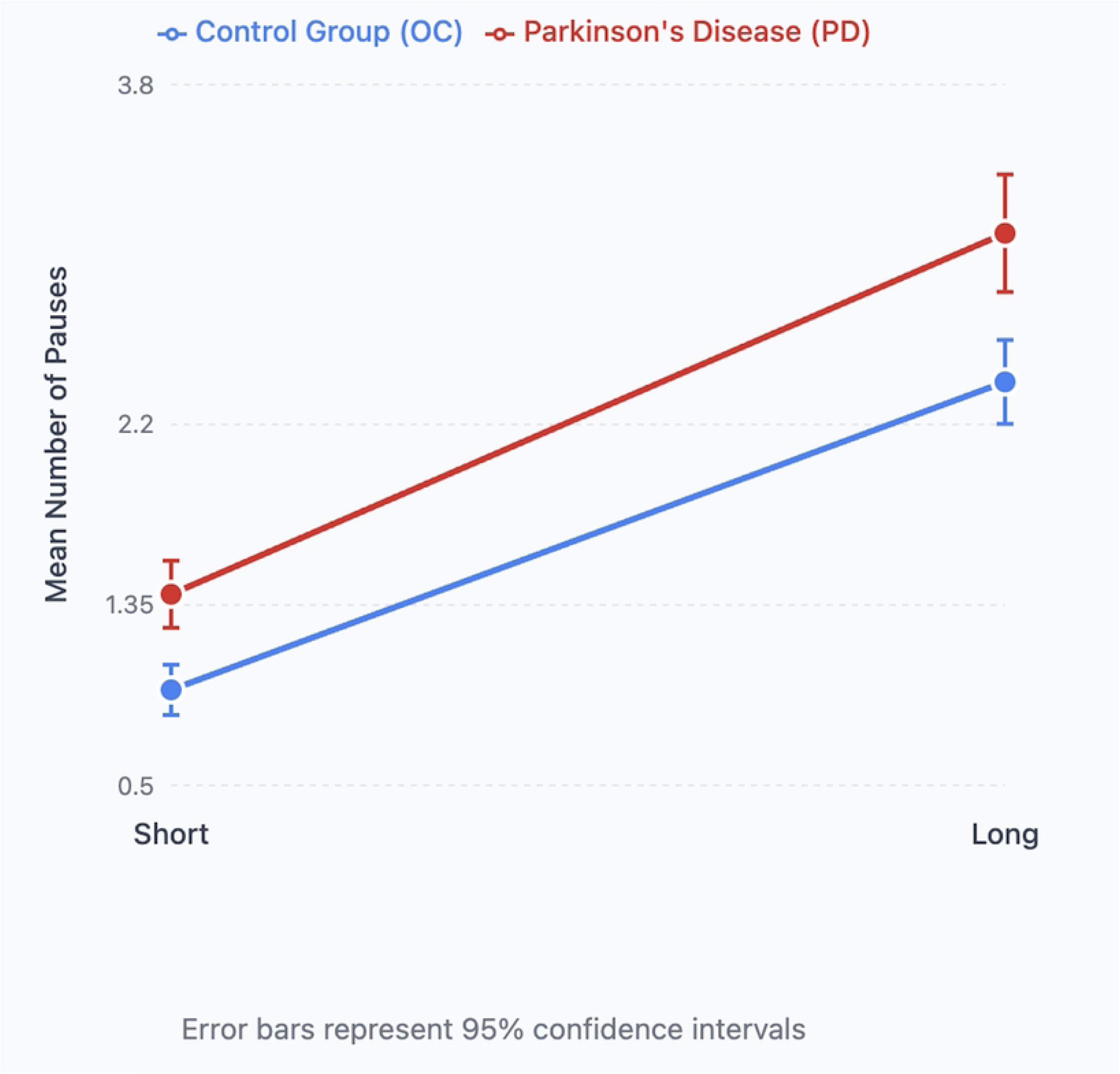
Pause numbers and sentence length. Mean number of pauses in short and long sentences in Parkinson’s Disease (PD) and Older Healthy (OC) participants. There was a significant main effect of disease status for the mean duration of pauses in both short and long sentences (*F*(1, 66) = 6.935, *p* = 0.011), with the PD group (mean = 0.396 seconds) producing longer pauses than the OC group (mean = 0.345 seconds). Additionally, a significant effect of sentence length was found (*F*(1, 66) = 67.873, *p* < 0.001), with longer sentences (mean = 0.411 seconds) displaying longer pause durations than shorter sentences (mean = 0.331 seconds). A significant interaction was found between disease status and the mean duration of pauses across sentence length (*F*(1, 66) = 4.263, *p* = 0.043). Fig 2 shows that the PD group had longer pause durations across both sentence types. It also highlights that longer sentences resulted in longer pauses than shorter sentences in both groups.

**Fig 2.**
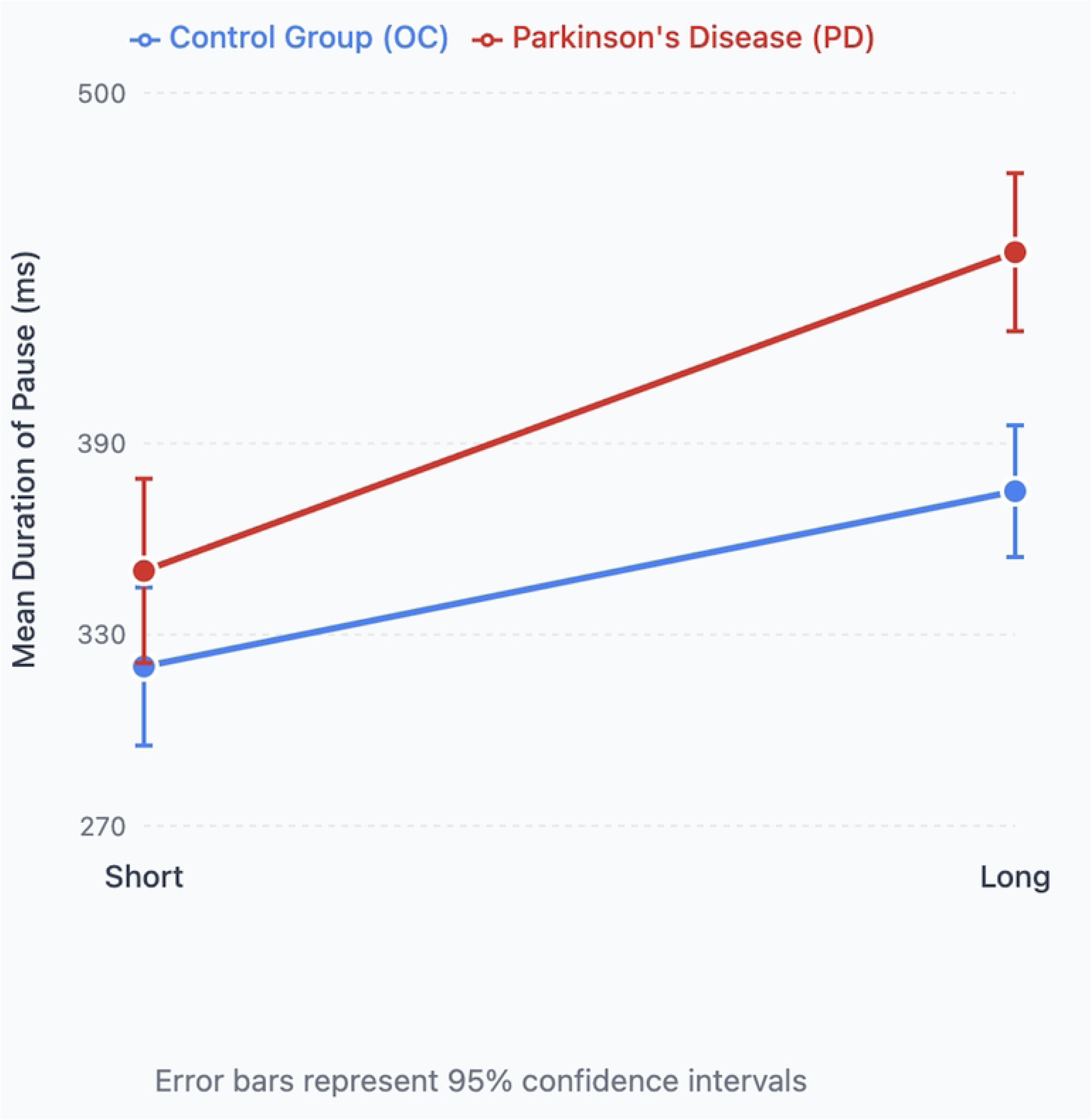
**Pause duration and sentence length**. Mean duration of pauses in short and long sentences in Parkinson’s Disease (PD) and Older Healthy (OC) participants. A significant main effect of disease status was found for the number of pauses produced in simple and complex sentences (F(1, 66) = 12.123, *p* < 0.001), with the PD group (mean = 1.938) producing more pauses than the OC group (mean = 1.404). Additionally, sentence complexity had a significant effect (*F*(1, 66) = 231.502, *p* < 0.001), with complex sentences (mean = 2.224) generating more pauses than simple sentences (mean = 1.118). A significant interaction was observed between the mean number of pauses and sentence complexity in relation to disease status (*F*(1, 66) = 6.604, *p* = 0.012). Fig 3 illustrates that the PD group had a greater number of pauses across both sentence complexity levels. It also demonstrates that more complex sentences yielded more pauses than simpler ones in both groups.

**Fig 3.**
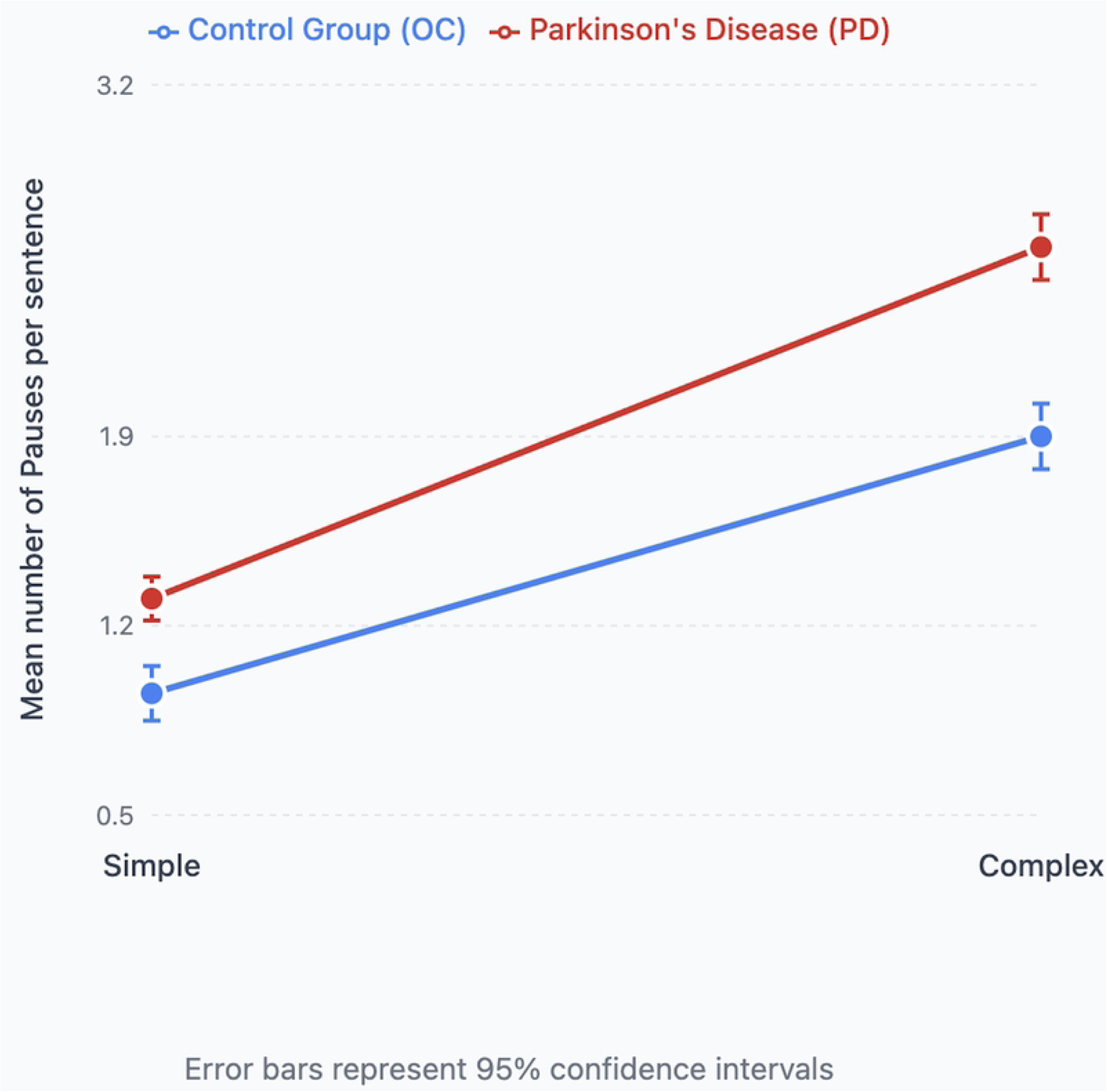
Pause numbers and sentence complexity. Mean number of pauses in simple and complex sentences in Parkinson’s Disease (PD) and Older Healthy (OC) participants. A significant main effect of disease status was found for the mean duration of pauses in simple and complex sentences (*F*(1, 66) = 9.456, *p* = 0.003), with the PD group (mean = 0.408 seconds) having longer pauses compared to the OC group (mean = 0.353 seconds). Additionally, sentence complexity had a significant effect on pause duration (*F*(1, 66) = 15.637, *p* < 0.001), with complex sentences (mean = 0.395 seconds) having longer pause durations than simple sentences (mean = 0.365 seconds). No significant interaction was found between sentence complexity and disease status for the mean duration of pauses (*F*(1, 66) = 0.890, *p* = 0.349). Fig 4 shows that the PD group had a higher mean duration of pauses than the OC group across both sentence types. It also demonstrates that complex sentences led to longer pauses than simple sentences in both groups.

**Fig 4.**
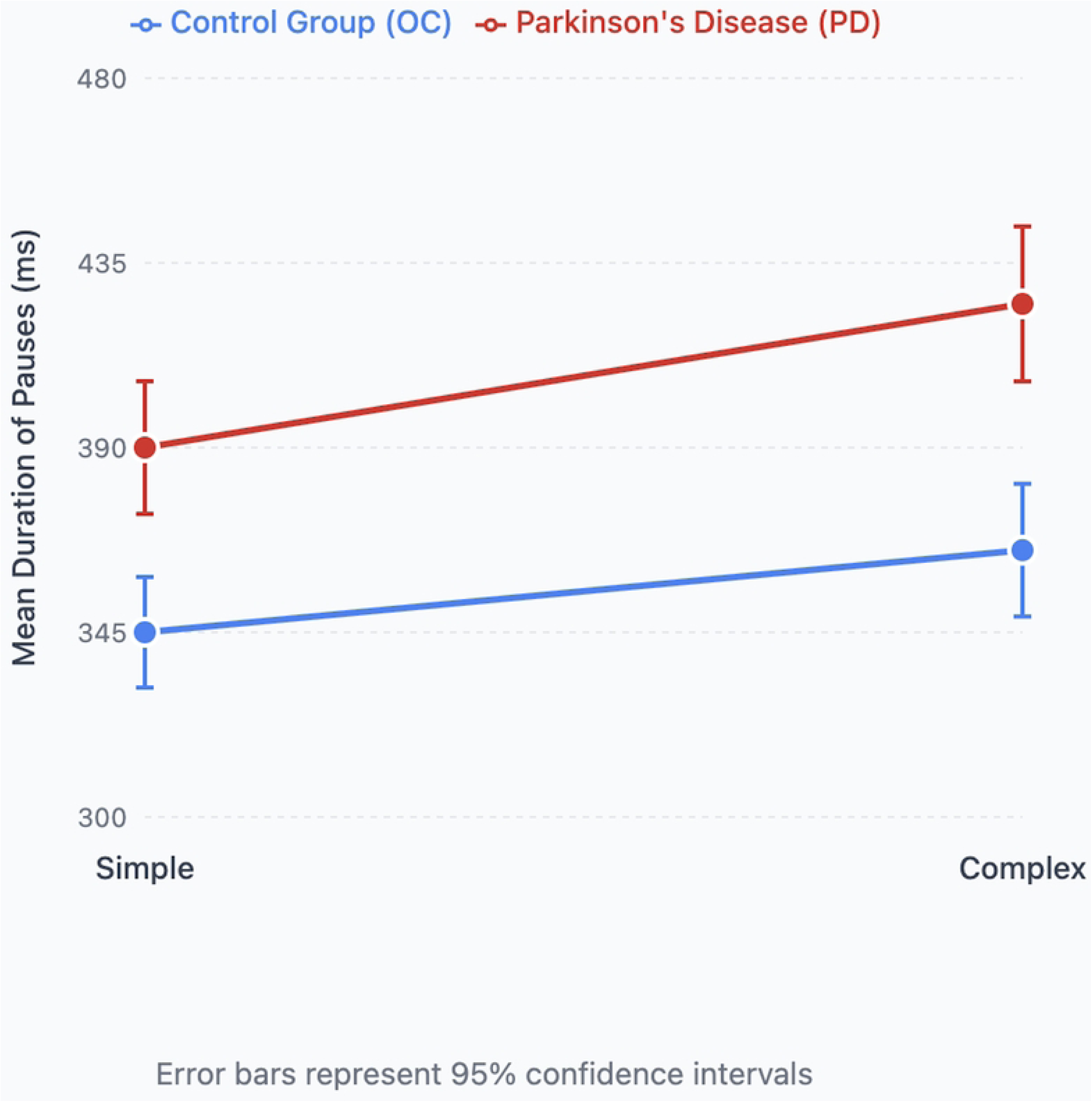
**Pause duration and sentence complexity**. Mean duration of pauses in milliseconds in simple and complex sentences in Parkinson’s Disease (PD) and Older Healthy (OC) participants.

#### Between-Group comparison

Significant differences between the PD and OC groups were found for several measures. For the mean duration of between-sentence pauses, the PD group had a significantly higher mean (mean = 0.960 seconds, Standard Deviation (SD) = 0.239) compared to the OC group (mean = 0.771 seconds, SD = 0.164), (*t*(49.144) = 3.691, *p* < 0.001). There was also a significant difference in the total number of pauses across the passage, with the PD group producing more pauses (mean = 38.77, SD = 15.119) than the OC group (mean = 28.05, SD = 9.997), (*t*(47.993) = 3.346, *p* = 0.002).

Further significant differences were observed for the mean number of pauses in short sentences, where the PD group had a higher mean (mean = 1.269, SD = 0.581) compared to the OC group (mean = 0.847, SD = 0.378), (*t*(47.469) = 3.450, *p* < 0.001). For long sentences, the PD group also produced more pauses (mean = 3.100, SD = 1.175) than the OC group (mean = 2.342, SD = 0.829), (*t*(66) = 3.116, *p* = 0.004). Additionally, for long sentence pauses, the PD group had a higher mean duration (mean = 0.446 seconds, SD = 0.117) compared to the OC group (mean = 0.375 seconds, SD = 0.059), (*t*(50.193) = 2.993, *p* = 0.004). The PD group also had more pauses in complex sentences (mean = 2.585, SD = 1.039) compared to the OC group (mean = 1.864, SD = 0.653), (*t*(66) = 3.497, *p* < 0.001).

No significant differences were observed between the groups for the following variables: mean duration of short pauses (*t*(38.338) = 1.329, *p* = 0.192), mean number of pauses in simple sentences (*t*(66) = 2.825, *p* = 0.006), mean duration of pauses in simple sentences (*t*(42.802) = 2.612, *p* = 0.018), and mean duration of pauses in complex sentences (*t*(41.504) = 2.854, *p* = 0.007).

### Research Question 2: Does sentence complexity and length affect pause behaviour?

#### Parkinson group (PD)

Significant differences were observed within the PD group for several comparisons with the adjusted α = 0.0125. There was a significant difference between the number of pauses in short and long sentences (*t*(29) = 13.060, *p* < 0.001), with long sentences (mean = 3.100, SD = 1.175) having more pauses than short sentences. Similarly, the duration of pauses in short and long sentences differed significantly (*t*(29) = 9.548, *p* < 0.001), with long sentences (mean = 0.446 seconds, SD = 0.117) having longer pause duration than short sentences (mean = 0.346 seconds, SD = 0.120). Additionally, the mean number of pauses was significantly higher in complex sentences (mean = 2.585, SD = 1.039) compared to simple sentences (mean = 1.292, SD = 0.581; *t*(29) = 5.394, *p* < 0.001). The duration of pauses was also significantly longer in complex sentences (mean = 0.426 seconds, SD = 0.109) compared to simple sentences (mean = 0.390 seconds, SD = 0.097; *t*(29) = 2.940, *p* = 0.006).

#### Older control group (OC)

In the OC group with the adjusted α = 0.0125, significant differences were observed between short and long sentences for the number of pauses (*t*(37) = 14.742, *p* < 0.001), with long sentences (mean = 2.342, SD = 0.829) having more pauses than short sentences (mean = 0.847, SD = 0.378). Similarly, the duration of pauses in short and long sentences was significantly different (*t*(37) = 12.641, *p* < 0.001), with long sentences (mean = 0.374 seconds, SD = 0.059) having longer pauses compared to short sentences (mean = 0.315 seconds, SD = 0.054). A significant difference was also found in the mean number of pauses between simple and complex sentences (*t*(37) = 6.542, *p* < 0.001), with complex sentences (mean = 1.864, SD = 0.653) having more pauses than simple sentences (mean = 0.944, SD = 0.434). However, no significant difference was found between the mean duration of pauses in simple and complex sentences (*t*(37) = 2.517, *p* = 0.016).

### Research Question 3: How do cognitive ability and dysarthria severity interact with sentence complexity and length to influence pause frequency and duration in PD?

#### Cognitive ability

Significant negative correlations were observed between cognitive ability, as measured by the MoCA, and the mean number of pauses in both short (*r_s_* (28) = −0.437, *p* = 0.016) and long sentences (*r_s_* (28) = −0.477, *p* = 0.008). Greater numbers of pauses were associated with lower cognitive ability. Similar negative correlations were found for the mean number of pauses in simple (*r_s_* (28) = −0.436, *p* = 0.016) and complex sentences (*r_s_* (28) = −0.492, *p* = 0.006), as well as the total number of pauses in the passage (*r_s_* (28) = −0.502, *p* = 0.005).

No significant correlations were found between cognitive ability and the duration of pauses in short sentences (*r_s_* (28) = −0.082, *p* = 0.667), long sentences (*r_s_* (28) = −0.196, *p* = 0.300), simple sentences (*r_s_* (28) = −0.051, *p* = 0.790), complex sentences (*r_s_* (28) = −0.187, *p* = 0.321), or between-sentence pauses (*r_s_* (28) = −0.273, *p* = 0.144).

#### Dysarthria severity

Significant positive correlations were found between dysarthria severity, as measured by the dysarthria rating, and the duration of pauses in short (*r_s_* (28) = 0.420, *p* = 0.021), long (*r_s_* (28) = 0.428, *p* = 0.018), and complex sentences (*r_s_* (28) = 0.509, *p* = 0.004). Longer pauses were associated with greater dysarthria severity. While not statistically significant, a borderline positive correlation was observed between dysarthria severity and the duration of pauses in simple sentences (*r_s_* (28) = 0.359, *p* = 0.052).

No significant correlations were found between dysarthria severity and the mean number of pauses in short (Spearman’s rho = 0.227, p = 0.227), long (Spearman’s rho = 0.035, p = 0.854), simple (Spearman’s rho = 0.055, p = 0.772), or complex sentences (Spearman’s rho = 0.188, p = 0.320), nor with the total number of pauses (Spearman’s rho = 0.114, p = 0.548).

## Discussion

### Overview

This study utilized a mixed design to investigate the influence of sentence complexity and sentence length on both the frequency and duration of silent pauses in speakers with Parkinson’s Disease (PD) compared to control speakers. It was hypothesized that PD speakers would exhibit an increased number and longer duration of pauses at all sentence levels—short, long, simple, and complex—relative to control speakers. The effect of sentence complexity was expected to influence both the frequency and duration of pauses, with more complex sentences leading to increased pause frequency and duration when compared to simple sentences. Similarly, sentence length was predicted to affect both the frequency and duration of pauses, with longer sentences expected to result in more frequent and prolonged pauses. Cognitive ability and dysarthria severity were anticipated to correlate with both the complexity and length of sentences; however, no specific predictions were made regarding these variables. The main findings from the study are as follows:

1. PD speakers exhibited a greater total number of pauses throughout the speech sample compared to control speakers.
2. PD speakers produced longer durations of between-sentence pauses relative to control speakers.
3. Significant differences between PD and control speakers were found at all sentence levels, except for the mean duration of pauses in short, simple, and complex sentences, as well as the mean number of pauses in simple sentences.
4. Within-group comparisons of sentence complexity revealed that both PD and control speakers produced more pauses in more complex sentences, though this effect was not observed in the duration of pauses across sentence complexities.
5. Sentence length had a significant effect within both groups, with longer sentences producing more pauses and longer durations of pauses in both PD and control groups.
6. Significant negative correlations were found between cognitive ability and the number of pauses in short, long, simple, and complex sentences, as well as the total number of pauses.
7. Significant positive correlations were observed between dysarthria severity and the duration of pauses in short, long, and complex sentences.

The results from this study demonstrate notable differences in the number and duration of silent pauses between PD and control speakers across all sentence levels. As hypothesized, PD speakers exhibited greater durations of between-sentence pauses, and at each sentence level, the frequency and duration of pauses were increased, except for the duration of pauses in short, simple, and complex sentences, as well as the mean number of pauses in simple sentences. The increased total number of pauses in PD speech aligns with prior research indicating that PD speakers require more preparation time to produce speech [48] and tend to produce atypical pauses within sentences [28]. This observation further supports the notion that PD speech is marked by reduced fluency, which may result from a combination of linguistic, motor, and cognitive factors, though the precise interaction between these factors remains unclear. Ash et al. [48] suggest that increased pauses may contribute to this reduced fluency, with pauses serving as indicators of cognitive and motor deficits in PD speech. Krivokapić [37] also suggests that sentence length may influence both pre- and post-boundary pause durations, which could explain the observed increase in between-sentence pauses. However, future studies need to independently investigate these factors to better understand their relative contributions.

Another potential explanation for the increased pause duration in PD speakers is the phenomenon of speech initiation hesitation. While this is unlikely to affect group differences, it may explain individual variability in pause duration. Moretti et al. [49] suggested that speech initiation delay, exacerbated by motor symptoms such as gait freezing in PD, may contribute to longer durations of between-sentence pauses. This should be considered when interpreting individual data.

Interestingly, no significant differences were found in the duration of pauses in short, simple, and complex sentences, nor in the mean number of pauses in simple sentences. This contrasts with some previous studies that have suggested PD speakers produce fewer pauses compared to control speakers, though the reasons for this discrepancy remain unclear. Stine-Morrow et al. [31] proposed that PD speakers might engage in “chunking” of speech, producing fewer but longer linguistic units in simpler sentences. This may account for the lack of statistical differences in simple sentences, although further research is needed to fully understand the linguistic units involved in PD speech.

The effects of sentence complexity and sentence length on the number and duration of pauses were consistent with predictions, with both factors contributing significantly to the differences observed between PD and control speakers. As expected, increased sentence complexity was associated with more within-sentence pauses, likely due to the greater cognitive load required to process more complex structures. For PD speakers, this cognitive demand likely results in the production of smaller linguistic units, leading to more frequent pauses [31], compounded by impaired physiological control of respiration and motor control [30]. Interestingly, while sentence complexity had a significant effect in both groups for the number of pauses, no significant effect was found for pause duration in the control group, which contradicts findings by Stine-Morrow et al. [31] that aging-related cognitive decline may have a similar effect to PD progression. This may be due to the use of a cognitive screening tool (MoCA), which indicated that all participants had intact cognitive abilities, potentially reducing the impact of age-related cognitive decline on the control group.

The effect of sentence length was as predicted, with both sentence constituent length (number of syllables) and higher information load contributing to an increase in both the number and duration of pauses. Previous studies have shown that longer sentences are associated with more frequent and prolonged atypical pauses [36,38]. However, this study did not control for the number of syllables, making it difficult to definitively attribute the observed effects to information load alone. The interaction of sentence length and syllable count likely requires further investigation to disentangle these contributing factors.

Significant negative correlations were found between cognitive ability and the number of pauses in several sentence categories (short, long, simple, complex, and total number of pauses). These results align with expectations, suggesting that cognitive ability influences the efficiency of lexical retrieval and word selection in PD speech [35]. Reduced cognitive resources in PD speakers likely lead to smaller linguistic units and an increase in pause frequency. However, no significant correlations were found between cognitive ability and the duration of pauses, which may be attributed to the fact that pause duration is influenced more by speech rate and articulatory movements, which are less affected by cognitive decline [31].

Disease severity was positively correlated with the duration of pauses in short, long, and complex sentences, with borderline significance in simple sentences. This finding aligns with previous research by Hammen and Yorkston [50], who observed that dysarthria severity is associated with increased pause duration, possibly due to dysarthric movements that delay speech initiation. This hypothesis warrants further investigation, particularly in terms of the relationship between dysarthria severity and speech rate, which was not controlled for in this study.

### Limitations and future directions

Several limitations should be considered when interpreting the results of this study. The lack of blinding in data analysis could introduce bias, as researchers were aware of group membership during analysis. Future studies could address this limitation by ensuring that data is anonymized until after analysis is completed. Additionally, the study did not control for the adaptation effect [51], which could have influenced the results. This limitation was particularly relevant for data collected during the COVID-19 pandemic, when resources for data collection had to be distributed in advance. Future research could mitigate this by conducting in-person sessions to better control for the adaptation effect.

Moreover, while Praat software was used to analyze pauses, adjustments to silence boundaries during analysis may have introduced inaccuracies in both temporal and speech initiation measurements. Despite these limitations, the inter-rater reliability was excellent, indicating that the results were consistent across individual datasets.

Finally, the use of Bonferroni corrections for statistical analyses increases the risk of type II errors, potentially leading to the exclusion of important variables. Future studies should explore alternative statistical methods to balance the risks of both type I and type II errors.

Future research could explore the use of silent pauses as potential biomarkers for disease progression in PD. Investigating whether pauses can serve as an outcome measure for speech therapy, particularly in assessing therapy efficacy, would be valuable. Additionally, research could examine whether silent pauses are consistent across different subtypes of dysarthria, contributing to the development of diagnostic screening tools.

## Conclusion

The findings of this study suggest that both cognitive demands and sentence complexity significantly influence the production of silent pauses in PD speech. Clinically, these findings indicate that silent pauses could serve as an objective measure of disease severity and treatment outcomes. Further research is needed to determine the potential utility of silent pauses as a tool for prioritizing patients within speech therapy services and assessing treatment efficacy. Additionally, understanding the theoretical underpinnings of these findings may lead to better therapeutic strategies for PD speakers.

## Acknowledgments

We would like to thank Dr. Arpita Bose and Dr. Christos Salis for their technical expertise and help with the data analysis for this project. Additionally, we would like to thank all the participants who took part in this study.

## Supporting information

S1 Table. Demographic information of Parkinson group

S2 Table. Demographic information of Control group

**S3 Table. Praat Coding Criteria.** based on DeDe & Salis (2019) and Reed (2020) - Only showing codes used in from initial criteria

S4 Table. Normality Test Results

## Appendices

**Appendix 1:** Criteria for Complexity of Sentence in Rainbow Passage (Based on Dede and Salis (2020)

| Simple | Complex |
| --- | --- |
| One independent clause containing a subject and a predicate | One or more dependent clauses to an independent clause |
| May contain compound nouns and verbs | Include at least one embedded clause |
| Can contain coordinating conjunctions (and, but, or) | Either have a subordinating clause (because, when, although) or a relative pronoun (that, which who etc.) |

Length of Sentence was determined by using the mean of the sentence length, with Long being above 17.21 words a sentence and short sentences below 17.21 words per sentence i.e. 327 words/19 sentences = 17.21 words per sentence

**Appendix 2:**
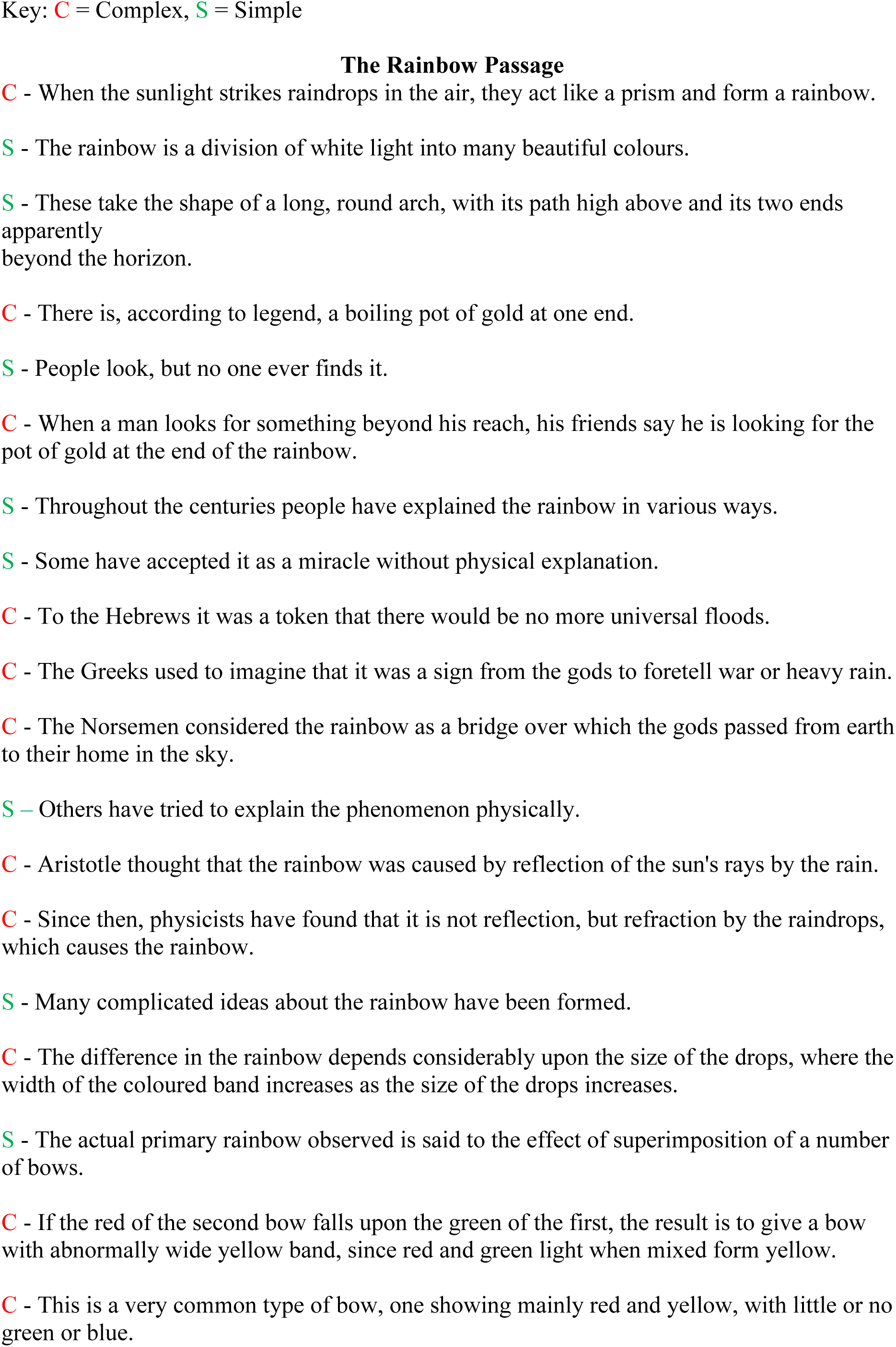
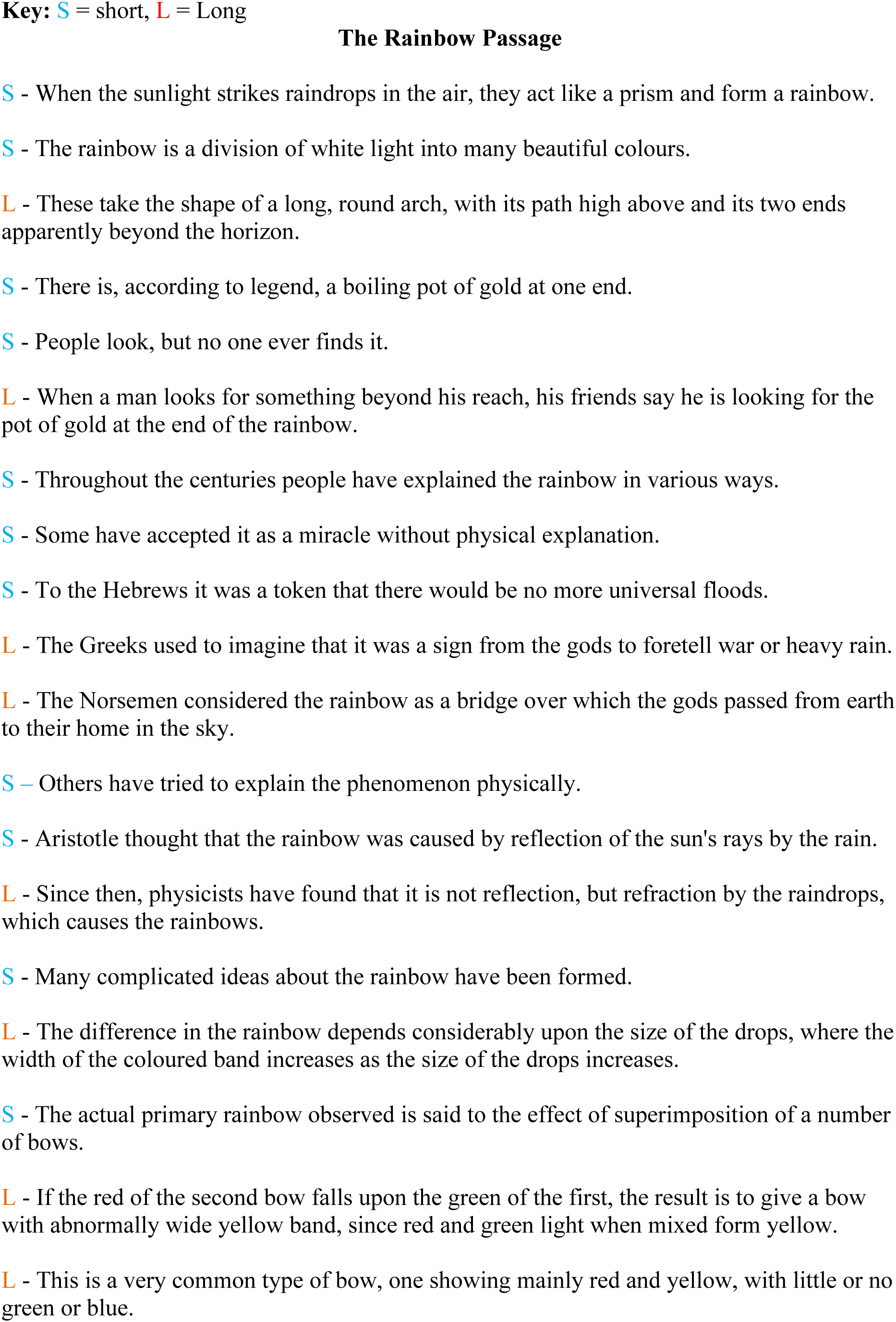
Categorisation of Sentences in Rainbow Passage

